# Enveloped viruses show increased propensity to cross-species transmission and zoonosis

**DOI:** 10.1101/2022.07.29.501861

**Authors:** Ana Valero-Rello, Rafael Sanjuán

## Abstract

The transmission of viruses between different host species is a major source of emerging diseases and is of particular concern in the case of zoonotic transmission from mammals to humans. Several zoonosis risk factors have been identified, but it is currently unclear which viral traits primarily influence this process, as previous work has focused on a few hundred viruses that are not representative of the actual viral diversity. Here we investigate fundamental virological traits that influence cross-species transmissibility and zoonotic propensity by interrogating a database of over 12,000 mammalian virus-host associations, obtained mainly from recent viral metagenomics projects. Our analysis reveals that enveloped viruses tend to infect more host species and are more likely to infect humans than non-enveloped viruses, while other viral traits such as genome composition, structure, size or the viral replication compartment play a minor role. This contrasts with the previous notion that viral envelopes did not significantly impact or even reduced zoonotic risk, and should help better prioritize outbreak prevention efforts. We suggest several mechanisms by which viral envelopes could promote cross-species transmissibility, including structural flexibility and evasion of viral entry barriers.

## Introduction

Viral cross-species transmission is at the origin of an increasing number of emerging diseases. This includes well-known zoonoses, such as HIV/AIDS, influenza, Zika, Ebola, monkey pox, and COVID, in addition to other animal and plant diseases. Understanding and ultimately predicting viral emergence has therefore become a major research goal^1–4^. Nearly 90% of known viral zoonoses originate from wild or domesticated mammals^5^. A number of risk factors have been identified, such as biodiversity loss^6^, species invasions^7^, wildlife trade^8^, viral host plasticity^9,10^, life history traits of reservoir hosts^11^, and host proximity to humans in terms of phylogenetic distance^12^ and interaction frequency^13^. Additionally, information about host usage and zoonotic propensity can be extracted from viral genomes by analyzing features such as codon or dinucleotide usage biases and their similarity to human gene transcripts^12,14,15^.

Despite these advances, it remains unclear how cross-species transmissibility and zoonosis depend on the fundamental properties of a virus. Previous work has emphasized that RNA viruses should be more prone to host jumps than DNA viruses owing to their extensive genetic diversity and fast adaptability^16,17^. Some recent studies have supported this view, whereas others have identified seemingly more relevant features, such as viral replication in the cytoplasm^18–20^. Some analyses have also suggested an effect of viral genome size or genome segmentation, whereas these findings have not been supported by others^9,13,18,21^.

A limitation of previous studies is that they rely essentially on well-known, ICTV-approved viruses, which constitute a small and non-representative subset of the mammalian virosphere. Importantly, viruses with socioeconomic implications, such as those causing human disease, have been preferentially investigated, critically biasing inferences on viral host usage, cross-species transmissibility, and zoonotic propensity^3^. Currently, though, most mammalian viruses are discovered through viral metagenomics^22^, including major initiatives such as PREDICT and the Global Virome Project^23^. This approach affords a still limited but less biased picture of viruses in nature, and has revealed a large number of new virus-host associations^24^. Viral metagenomics often provides only one or a few short sequences of each new virus. Although this precludes certain analyses, these sequences inform about fundamental viral features, such as the nature, size, and structure of their genetic material. It is therefore possible to leverage this information to examine how such features influence the ability of viruses to infect multiple host species and cause zoonoses.

## Dataset

We extracted from the VIRION database 5149 viruses belonging to 36 families and 1599 hosts species from 20 orders, comprising in total 12,888 virus-host associations (**Dataset S1**). All viruses were ratified by the NCBI taxonomy site and were assigned a viral family. Most (77.8%) were not ICTV-approved viruses and thus may be genuine species, higher-order taxa, or viral subtypes. Approximately half of the viruses (52.6%) had a single sequence record in NCBI Entrez, whereas some had thousands (**Fig. 1a**). As expected, the probability that a virus was found in multiple hosts depended strongly on the number of sequence records available for that virus, N (**Fig. 1b, 1c**). We therefore included N as a covariate in subsequent analyses.

**Figure 1.**
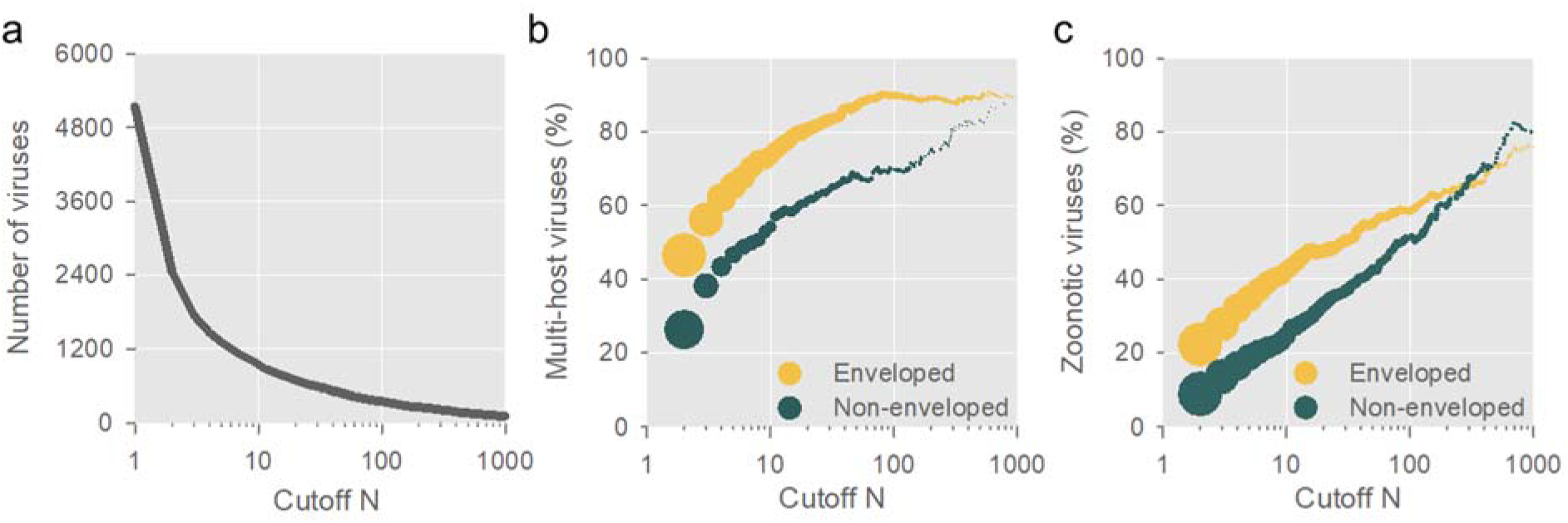
Viral discovery trends. **a**. Most viruses are discovered through metagenomics and have a very small number of sequence records. The x-axis shows the minimal number of sequences required for inclusion in our analysis (cutoff N), and the number of viruses meeting this condition is shown on the y-axis. **b**. The fraction of multi-host viruses increases more rapidly with the cutoff N for enveloped viruses than for non-enveloped viruses. Dot sizes are proportional to the number of viruses included in each cutoff. The dots are shown for N ≥ 2, N ≥ 3, and so on. Dots for N ≥ 1 were too large for visualization and are omitted. **c**. A similar trend was found for zoonotic viruses. For the minute fraction of viruses with many available sequences, non-enveloped are similarly or even more zoonotic than enveloped viruses, reproducing previous findings (see text).

We considered five dichotomous and one continuous variables defining fundamental viral features: whether the genome is made of RNA or DNA, single or double stranded, segmented or non-segmented, whether the virus replicates in the cytoplasm or the nucleus, whether the virus is enveloped or non-enveloped, and the viral genome size. These features are conserved within viral families and hence their imputation was straightforward for all viruses, even if only few short sequences were available without further characterization.

### Cross-species transmissibility

Since zoonosis is a special case of viral cross-species transmission, we first examined whether or not viruses were found in multiple mammalian host species (multi-host viruses), excluding humans to reduce bias. An exploratory analysis revealed that the fraction of multi-host viruses increased more rapidly with N for enveloped viruses than for non-enveloped viruses (**Fig. 1b**). This difference was still obvious when we combined the envelope variable with each of the other dichotomous viral traits considered (**Fig. S1**). For instance, the fraction of multi-host viruses was higher for enveloped DNA viruses than for non-enveloped RNA viruses.

To formally assess the effects of different viral features on the probability of infecting multiple non-human host species, we performed a binary logistic regression analysis. For this, we initially set a cutoff number of sequences N ≥ 5 to avoid poorly explored or ill-defined viruses. This condition was met for 1305 viruses, of which nearly half (42.4%) were not ICTV-approved. We found a strongly significant effect of the viral envelope on the multi-host probability (*P* < 0.0001), whereas all other factors were non-significant, except for a marginally significant increase associated with single-stranded genomes (**Table S1**; **Fig. 2a**). As suggested previously^21^, viruses with small genomes tended to be less often found in multiple hosts, but this effect was accounted for by the fact that these are typically non-enveloped viruses.

**Figure 2.**
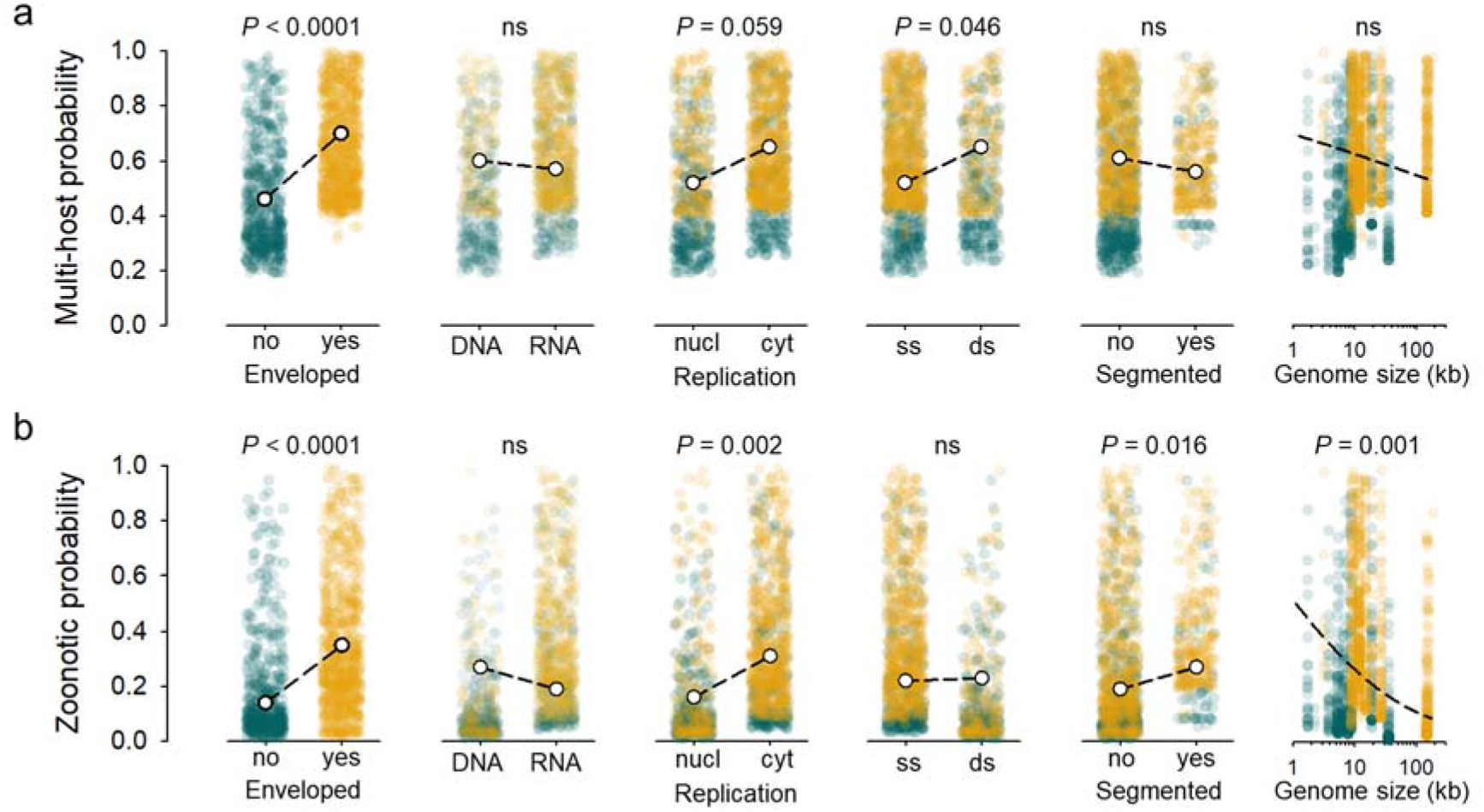
Binary logistic regression analysis for viruses with N ≥ 5 available sequences. **a**. Scatter plots show the predicted probability of being a multi-host virus for each of the 1305 viruses included in this analysis. Orange dots indicate enveloped viruses and green dots non-enveloped viruses. White dots and dashed lines indicate the marginal mean predicted by the model. P-values for each predictor variable are shown. N was included as a covariate in the analysis but is not shown. A summary of the model statistics is provided in **Table S1. b**. Same plots for zoonosis probability. A summary of the model statistics is provided in **Table S2**.

We checked that differences between enveloped and non-enveloped viruses were not driven by specific viral families (**Fig. 3**; **Fig. S2**). Indeed, when each viral family was considered as a categorical factor in a stepwise logistic regression, having an envelope was again the only viral feature significantly associated with multi-host probability (Wald test: *P* < 0.0001). We also verified that our conclusions were robust to the cutoff N used by performing the logistic regression with all 5149 viruses, those with N ≥ 2, N≥ 3, and so on for all cutoff values until the sample size became too small to detect any effect (**Fig. S3**).

**Figure 3.**
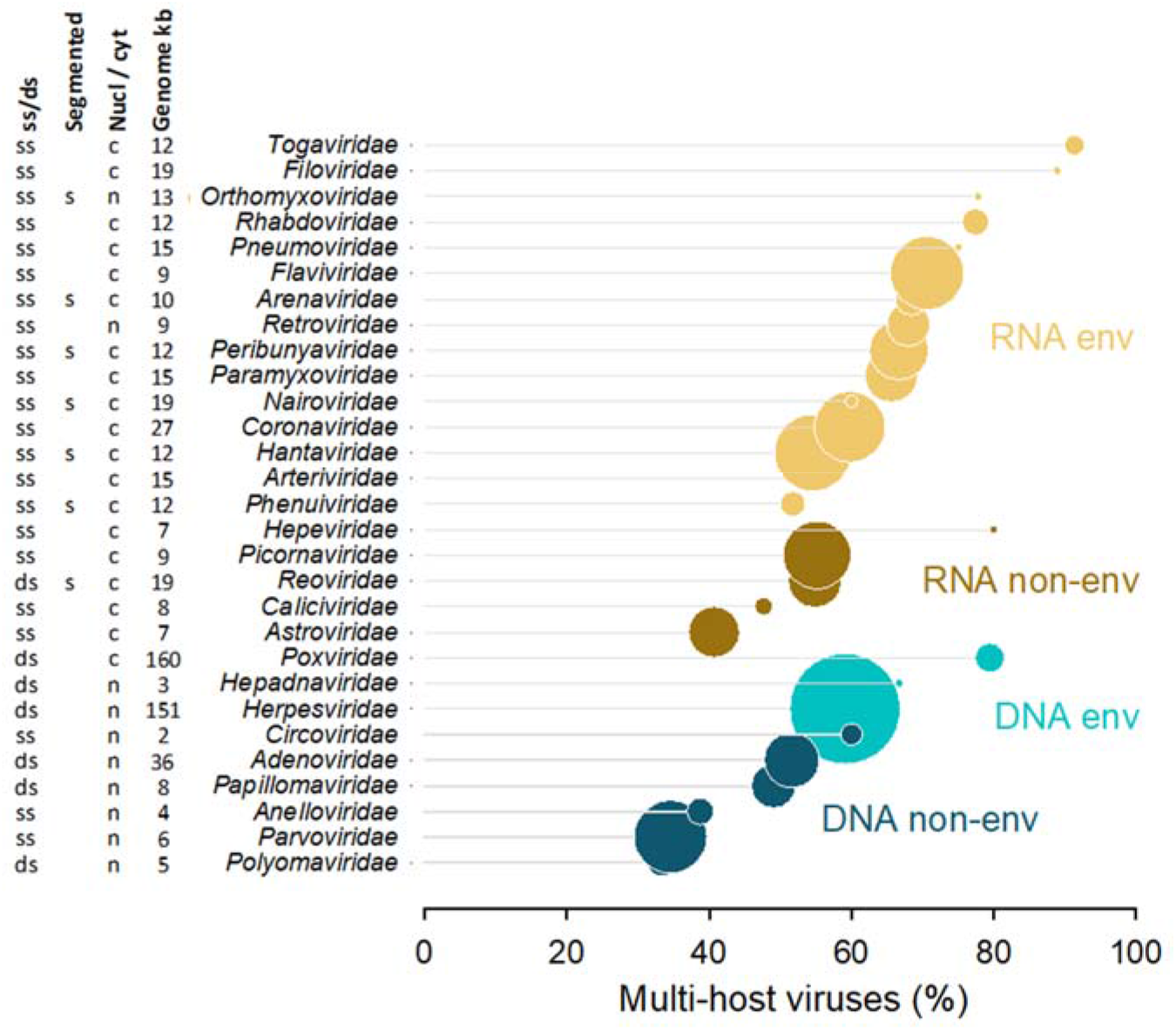
Fraction of multi-host viruses by viral family. Families with five or more different viruses, each having N ≥ 5 sequence records are shown. The size of the bubbles is proportional to the number of viruses in the family. The largest bubble corresponds to *Herpesviridae*, which has 127 different viruses with N ≥ 5, whereas the smallest corresponds to *Hepeviridae*, with 5 viruses. The presence of an envelope and the type of nucleic acids are color-coded. On the left are shown the other four viral features considered in our analyses. See **Fig. S2** for a plot that also considers the influence of N on the multi-host fraction.

We also performed an alternative analysis in which we examined the host species count per virus (host breadth). For this, we implemented a negative binomial regression, which confirmed that having an envelope was the viral feature most significantly associated cross-species transmissibility (*P* < 0.0001; **Table S2**). Host breadth contained more information than the binary multi-host variable, and the regression explained a higher fraction of the total deviance (53.4%). However, this approach was probably less robust to biases in the dataset, and the model fit was complicated by the strong overdispersion shown by host breadth (**Fig. S4**).

### Zoonotic viruses

We then focused on whether or not viruses were zoonotic, i.e. capable of infecting humans. As above, the fraction of zoonotic viruses increased more rapidly with N for enveloped viruses than for non-enveloped viruses (**Fig. 1c**). A binary logistic regression analysis for viruses with N ≥ 5 sequence records confirmed a strongly significant effect of the viral envelope, with an estimated 2.5-fold increase in zoonotic propensity compared to non-enveloped viruses (**Fig. 2b; Table S3**). Statistical significance was robust to the cutoff N used, as evidenced when we considered all 5419 viruses in the dataset, those with N ≥ 2, N≥ 3, and up to N ≥ 65 (**Fig. S3**). In addition, we found that viruses replicating in the cytoplasm were 1.9 times more likely to be zoonotic than those replicating in the nucleus, in accordance with previous studies^18,19^. We also detected a slightly higher propensity of segmented viruses to zoonosis compared to non-segmented viruses, also consistent with previous suggestions^13^, and a decreasing zoonotic probability for viruses with larger genomes. These effects were weaker or not observed when we examined the cross-species transmissibility of viruses excluding humans, suggesting human-specific factors or, more likely, biases in human-infective virus datasets.

## Discussion

Our results contrast with previous work suggesting that enveloped viruses were similarly or even less prone to zoonosis than non-enveloped viruses, and that the main viral feature associated with zoonosis was replication in the cytoplasm^9,13,18,21^. This is probably due to the fact that previous analyses focused only on approximately 350 highly-studied viruses. Supporting this, whether viruses replicate in the cytoplasm was the only statistically significant feature explaining zoonotic propensity when our logistic regression analysis was performed with the 353 viruses that had N ≥ 100 available sequence records, reproducing previous findings^18^. In addition to human-centric bias, a possible caveat with the effect of cytoplasmic replication on zoonotic risk is that this feature is typical of RNA viruses (93.1% and 96.5% coincidence for ICTV viruses and our dataset, respectively), making it difficult to separate the contribution of these two traits. We therefore suggest caution concerning whether RNA genomes or cytoplasmic replication is a factor driving zoonotic propensity.

Previous work has also suggested that enveloped viruses tend to be less transmissible among humans than non-enveloped viruses, potentially due to their lower stability in the environment^25^. We suggest that this observation can be viewed as a consequence of the broader host range exhibited by enveloped viruses. Hosts harbors a mixture of ancient vertically inherited and recent horizontally transmitted viruses^26^. According to our results, enveloped viruses cause spillover infections more frequently than non-enveloped viruses. Although spillovers are frequent, only a small fraction of these events result in successful horizontal transmissions. Therefore, the fraction of human-infective viruses that do not achieve successful human-human transmission should be enriched in enveloped viruses.

A recent work ranked 889 wildlife animal viruses according to the risk of animal-to-human spillover, based on a systematic analysis of expert opinion^13^. Interestingly, despite the viral envelope was given a negative score in this analysis, 47 of the 50 top-listed viruses were enveloped, although these represented only about half of the total viruses considered. This emphasizes how the importance of enveloped viruses has been previously overlooked, despite the fact that the vast majority of remarkable zoonotic viruses in human history are enveloped, including poxviruses (e.g smallpox, monkey pox), morbilliviruses (e.g. measles), rhabdoviruses (e.g. rabies), coronaviruses (e.g. COVID), flaviviruses (e.g. Zika), orthomyxoviruses (e.g. influenza), and retroviruses (e.g. HIV).

The mechanistic basis for the broader host tropism displayed by enveloped viruses remains to be investigated. Receptor-mediated viral entry is a critical stage in viral infection and cross-species transmission^27^. In principle, envelope proteins should be structurally less constrained than capsid proteins, which are more rigid. This might allow enveloped viruses to bind cellular receptors from different host species in a more flexible manner, to bind a greater number of alternative receptors, or to better accommodate host-switch mutations without compromising other functions. In addition, enveloped viruses can enter host cells through apoptotic mimicry, a process by which viral particles are engulfed by cells camouflaged as phosphatidylserine-rich apoptotic bodies^28^. This process has been shown to be relevant for different enveloped viruses, such as alphaviruses, ebolaviruses and poxviruses, all of which display a broad host range. Finally, it is also possible that envelopes contribute to cross-species transmissibility by helping viruses evade host immunity^29^. Future work might elucidate whether these or other processes drive the increased ability of enveloped viruses to infect different hosts and cause zoonoses.

## Methods

### Dataset and data curation

Viruses associated to mammalian hosts were obtained from the VIRION database^24^ (www.viralemergence.org/virion), last accessed in July 2022. Duplicates by means of serology, molecular or isolation were removed, resulting in unique virus-host associations. Viruses that were not classified at the family level, not resolved NCBI taxonomy, with missing names or named after viral family only [-viridae sp.]) were filtered out. The total counts of nucleotide sequences available for each virus (including any lower taxonomic level) were obtained from NCBI, and viruses without nucleotide records were removed. The vast majority of viruses with a single sequence record originated from a single report and were associated with a single host species, although 1.2% were multi-host viruses because evidence of infection was obtained by other methods (serological, PCR, etc). Hosts not resolved at the species level or with uncertainty in their identification according to the VIRION database (tagged as HostFlagID=TRUE, not binomial scientific name) were also removed. Human-exclusive viruses were not included. Further manual curation was done to clear isolates suspected to infect only non-mammalian hosts according to literature, such as viruses belonging to the family *Picobirnaviridae*, which were initially believed to infect animals but were later suggested to be bacteriophages^30^. The family *Smacoviridae* was removed due to its poor characterization, which did not allow assigning all features unambiguously.

### Viral features

Since the sequence information available for each virus was sufficient for taxonomical classification at the family level, fundamental features such as the genetic material (RNA/DNA), the presence of an envelope, the replication site (cytoplasmic/nuclear), genome strands (single/double), genome segmentation, and genome size could be assigned to each virus, and were obtained from either ViralZone or ICTV. For genome size, we took the value corresponding to a prototypical member of the family (**Table S4**).

### Statistical analysis

The total count of host species was obtained for each virus (host breadth), from which the binary response variable multi-host was calculated. Zoonotic viruses were defined as for those infecting humans and at least one additional mammal species. Binary predictors were DNA/RNA genome, enveloped/non-enveloped, nuclear/cytoplasmic replication, single/double stranded, and segmented/non-segmented. The quantitative predictors N and genome size were log-transformed. Generalized linear models (GLMs) using different distribution families and link functions were benchmarked (**Table S5**). For viral host breadth, negative binomial models were judged more convenient than Poisson models due to data overdispersion (**Fig. S4**). For these models, square root and log link functions performed similarly, but we favored the canonical link function log. For the binary response variables multi-host and zoonotic, we used binomial regression models, and we also selected the canonical link function log, making the model equivalent to a binary logistic regression. The selected link function performed similarly well as the clog-log and square root functions. We verified that, in all examined models, the presence/absence of an envelope was the viral trait that explained each of the three response variables with the highest significance (*P* < 0.0001 in all cases). We also explored generalized additive models (GAMs), but these did not provide an obvious improvement in performance over GLMs and had the drawback of being less interpretable. For the binary logistic regression on multi-host probability that incorporated the viral family as a predictor, the 29 families having at least five viral species were included. Each was converted to a binary variable, and the logistic regression was performed in a stepwise forward manner, such that each new predictor was added or not depending on whether a significant increase in the model likelihood was achieved, according to a Wald test. Statistical analyses were performed with R functions glm, glm. nb (R package MASS), and gam (R package mgcv), and SPSS v22.

## Acknowledgments

We thank Jérémy Dufloo, Pilar Domingo-Calap, and Iván Andreu-Moreno for helpful comments. This work was financially supported by Advanced Grant 101019724 - EVADER and grant PID2020- 118602RB-I00 – ZooVir from the Spanish Ministerio de Ciencia e Innovación to to R.S.

