## Supplemental tables and figures for "Enveloped viruses show increased propensity to cross-species transmission and zoonosis"

### Supplementary Tables and Figures

**Table S1.** Binary logistic regression analysis examining the probability that a virus is found in multiple hosts (excluding humans) as a function of the indicated features, focusing on the 1305 viruses with $N \geq 5$ available sequences.

| **Feature** | **Class** | **Number of viruses** | **Marginal mean** | **Likelihood ratio chi-square** | **p-value** |
|---|---|---|---|---|---|
| Enveloped | Yes | 788 | 0.70 ± 0.03 | 26.94 | <0.0001 |
| | No | 517 | 0.46 ± 0.03 | | |
| Genetic material | RNA | 843 | 0.57 ± 0.04 | 0.097 | 0.755 |
| | DNA | 462 | 0.60 ± 0.04 | | |
| Replication site | Cytoplasm | 816 | 0.65 ± 0.04 | 3.571 | 0.059 |
| | Nucleus | 489 | 0.52 ± 0.04 | | |
| Genome strand | Single | 924 | 0.52 ± 0.04 | 3.979 | 0.046 |
| | Double | 381 | 0.65 ± 0.04 | | |
| Segmented | Yes | 311 | 0.56 ± 0.04 | 1.75 | 0.186 |
| | No | 994 | 0.61 ± 0.02 | | |
| $\log_{10}$(genome_size) | | Coeff: | -0.32 ± 0.25 | 1.607 | 0.205 |
| $\log_{10}$(N) | | Coeff: | 1.12 ± 0.09 | 188.686 | <0.0001 |
| Intercept | | | | 17.02 | <0.0001 |

**Table S2.** Negative binomial regression analysis examining host breadth (number of host species excluding humans) as a function of the indicated features, focusing on the 1305 viruses with N ≥ 5 sequence records.

| **Feature** | **Class** | **Number of viruses** | **Marginal mean** | **Likelihood ratio chi-square** | **p-value** |
|---|---|---|---|---|---|
| Enveloped | Yes | 788 | 4.83 ± 0.32 | 24.48 | <0.0001 |
| | No | 517 | 2.95 ± 0.20 | | |
| Genetic material | RNA | 843 | 3.71 ± 0.30 | 0.052 | 0.820 |
| | DNA | 462 | 3.84 ± 0.35 | | |
| Replication site | Cytoplasm | 816 | 4.42 ± 0.33 | 6.15 | 0.013 |
| | Nucleus | 489 | 3.23 ± 0.25 | | |
| Genome strand | Single | 924 | 3.53 ± 0.27 | 0.890 | 0.346 |
| | Double | 381 | 4.03 ± 0.36 | | |
| Segmented | Yes | 311 | 3.56 ± 0.27 | 1.88 | 0.170 |
| | No | 994 | 4.01 ± 0.17 | | |
| $log_{10}$(genome_size) | | Coeff: | -0.06± 0.13 | 0.174 | 0.677 |
| $log_{10}$(N) | | Coeff: | 0.91 ± 0.04 | 804.44 | <0.0001 |
| Intercept | | | | 1120.08 | <0.0001 |

**Table S3.** Binary logistic regression analysis examining the probability that a virus is zoonotic as a function of the indicated features, focusing on the 1305 viruses with $N \geq 5$ available sequences.

| **Feature** | **Class** | **Number of viruses** | **Marginal mean** | **Likelihood ratio chi-square** | **p-value** |
|---|---|---|---|---|---|
| Enveloped | Yes | 788 | 0.35 ± 0.03 | 28.332 | <0.0001 |
| | No | 517 | 0.14 ± 0.02 | | |
| Genetic material | RNA | 843 | 0.19 ± 0.03 | 1.677 | 0.195 |
| | DNA | 462 | 0.27 ± 0.04 | | |
| Replication site | Cytoplasm | 816 | 0.31 ± 0.04 | 10.065 | 0.002 |
| | Nucleus | 489 | 0.16 ± 0.03 | | |
| Genome strand | Single | 924 | 0.22 ± 0.03 | 0.113 | 0.737 |
| | Double | 381 | 0.23 ± 0.04 | | |
| Segmented | Yes | 311 | 0.27 ± 0.03 | 5.808 | 0.016 |
| | No | 994 | 0.19 ± 0.02 | | |
| $\log_{10}$(genome_size) | | Coeff: | -1.07 ± 0.32 | 11.01 | 0.001 |
| $\log_{10}$(N) | | Coeff: | 1.34 ± 0.09 | 279.945 | <0.0001 |
| Intercept | | | | 170.821 | <0.0001 |

**Table S4.** Viruses used to impute genome sizes.

| Viral family | Reference virus for genome size |
|---|---|
| *Adenoviridae* | Human adenovirus C |
| *Anelloviridae* | Torque teno virus 1 |
| *Arenaviridae* | Lymphocytic choriomeningitis virus |
| *Arteriviridae* | Porcine reproductive and respiratory |
| *Asfarviridae* | African swine fever virus |
| *Astroviridae* | Human astrovirus 1 |
| *Bornaviridae* | Borna disease virus |
| *Caliciviridae* | Norwalk virus |
| *Circoviridae* | Porcine circovirus 1 |
| *Coronaviridae* | Human coronavirus 229E |
| *Filoviridae* | Zaire ebolavirus |
| *Flaviviridae* | Hepatitis C virus |
| *Genomoviridae* | Human associated gemyvongvirus 1 |
| *Hantaviridae* | Hantaan virus |
| *Hepadnaviridae* | Hepatitis B virus |
| *Hepeviridae* | Hepatitis E virus |
| *Herpesviridae* | Human alphaherpesvirus 1 |
| *Matonaviridae* | Rubella virus |
| *Nairoviridae* | Crimean-Congo hemorrhagic fever virus |
| *Nodaviridae* | Nodamura virus |
| *Orthomyxoviridae* | Influenza A virus (H1N1) |
| *Papillomaviridae* | Human papillomavirus type 16 |
| *Paramyxoviridae* | Mumps virus |
| *Parvoviridae* | Bovine parvovirus |
| *Peribunyaviridae* | Bunyamwera virus |
| *Phenuiviridae* | Rift Valley fever virus |
| *Picornaviridae* | Equine rhinitis B virus 1 |
| *Pneumoviridae* | Human respiratory syncytial virus |
| *Polyomaviridae* | Simian virus 40 |
| *Poxviridae* | Rabbit fibroma virus |
| *Reoviridae* | Bluetongue virus |
| *Retroviridae* | Human immunodeficiency virus 1 |
| *Rhabdoviridae* | Rabies virus |
| *Tobaniviridae* | Human torovirus |
| *Togaviridae* | Chikungunya virus |

**Table S5.** Benchmarking of different generalized linear models for the analysis of viral host breadth, multi-host status, and zoonotic status.

| Response variable | Type | Model | Link function[a] | Goodness of fit[c] | | |
|---|---|---|---|---|---|---|
| | | | | Pseudo $r^2$ | AIC | RMSE |
| Host breadth | Count | Negative binomial | log | 0.5258 | 6909.44 | 15.525 |
| | Count | Negative binomial | square root | 0.5147 | 6923.82 | 11.753 |
| | Count | Negative binomial | identity | 0.4731 | 6977.89 | 12.331 |
| | Count | Poisson | log | 0.4804 | 12375.89 | 4.789 |
| | Count | Poisson | square root | 0.4847 | 12304.53 | 5.209 |
| | Count | Poisson | identity | 0.4287 | 13225.65 | 5.583 |
| Multi-host | Binary | Binomial | logit | 0.1900 | 1232.93 | 0.448 |
| | Binary | Binomial | probit | 0.1871 | 1236.65 | 0.449 |
| | Binary | Binomial | cloglog | 0.1915 | 1230.99 | 0.448 |
| Zoonotic | Binary | Binomial | logit | 0.2765 | 1065.82 | 0.387 |
| | Binary | Binomial | probit | 0.2789 | 1062.65 | 0.387 |
| | Binary | Binomial | cloglog | 0.2835 | 1056.53 | 0.386 |

[a]The logarithm transform (log) is the canonical link function for the negative binomial, Poisson, and binomial regression models. cloglog: complementary log-log function.

[b]Pseudo $r^2$: deviance explained by the model, i.e. the relative reduction in deviance compared to an intercept-only model; AIC: Akaike information criterion, RMSE: root mean square error.

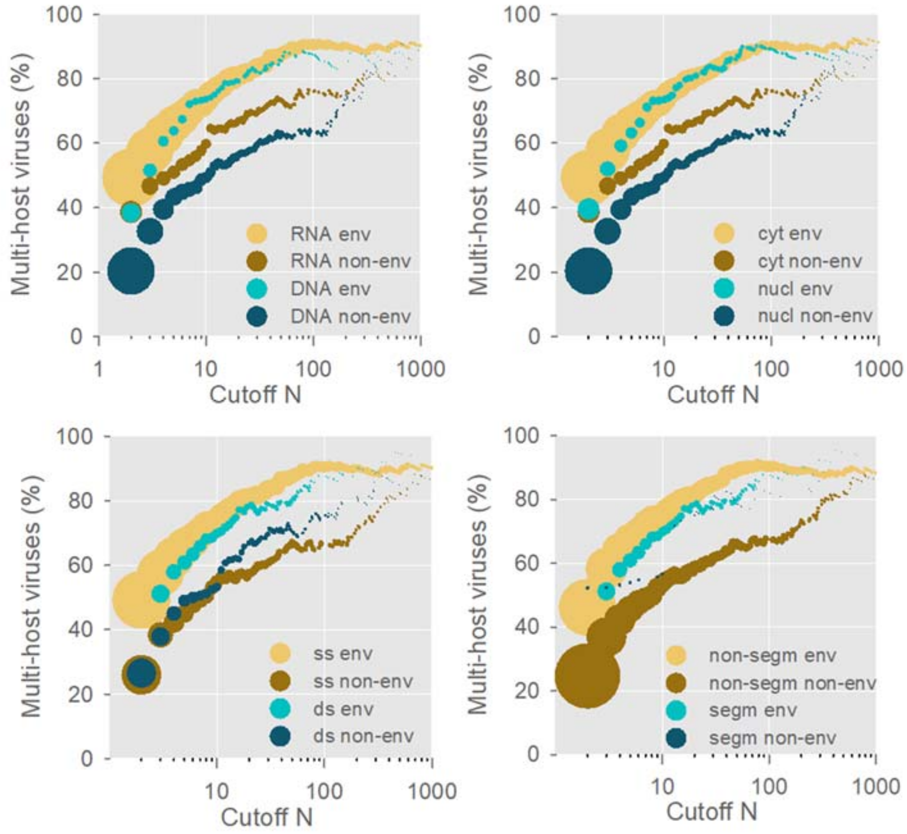


**Figure S1. Differences between enveloped and non-enveloped viruses considering other viral features.** The fraction of multi-host viruses is shown for each cutoff N. In each panel, the data are stratified by the presence of an envelope and another viral trait (DNA/RNA, cytoplasmic/nuclear replication, single/double-stranded, genome segmentation). Dot sizes are proportional to the number of viruses included in each cutoff. The dots are shown for $N \geq 2$, $N \geq 3$, and so on. Dots for $N \geq 1$ were too large for visualization and are omitted.

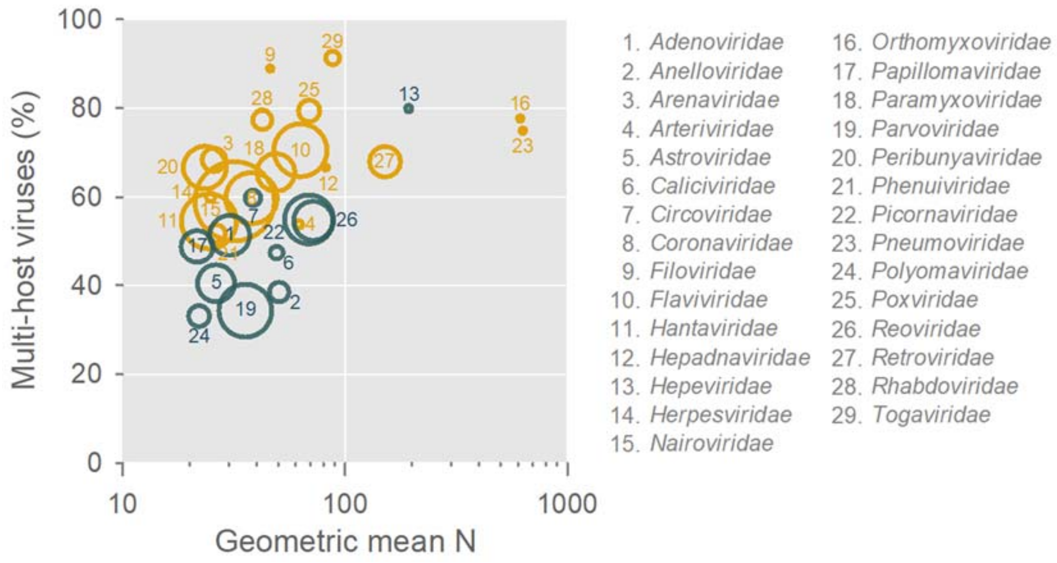


**Figure S2. Fraction of multi-host viruses by viral family as a function of N.** The 29 families with five or more different viruses, each having N ≥ 5 available sequences are shown. The size of the bubbles is proportional to the number of viruses in each family, identified by numbers. Enveloped viruses are colored in yellow, and non-enveloped viruses in dark green.

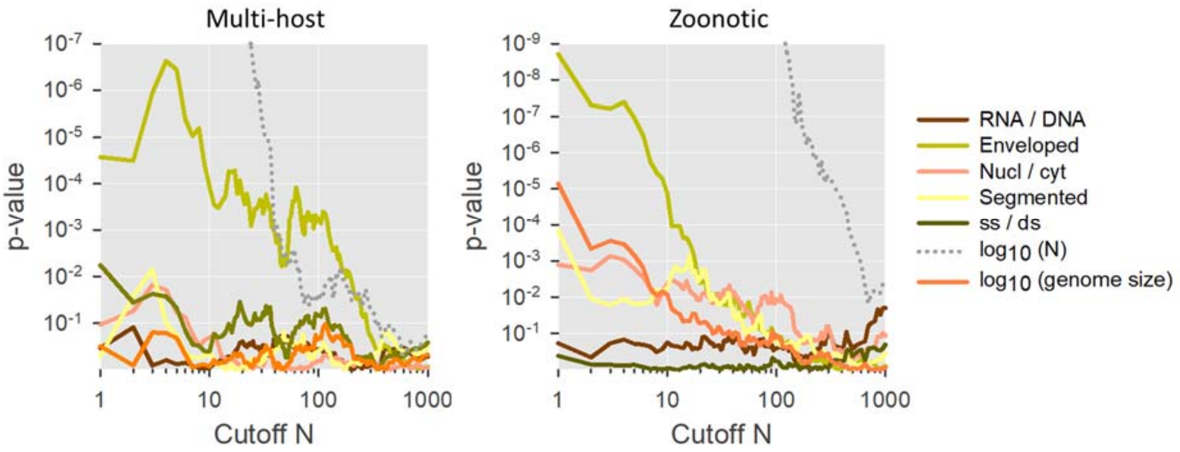


**Figure S3. Results of the binary logistic regression for different cutoff N values.** The y-axis shows the p-value obtained for each of the explanatory variables. The analysis was run for viruses with N ≥ 1, 2, …, 1000 sequences. The p-values for the covariate log(N) are shown out of scale to help visualize relevant p-values. Left: multi-host viruses; right: zoonotic viruses.

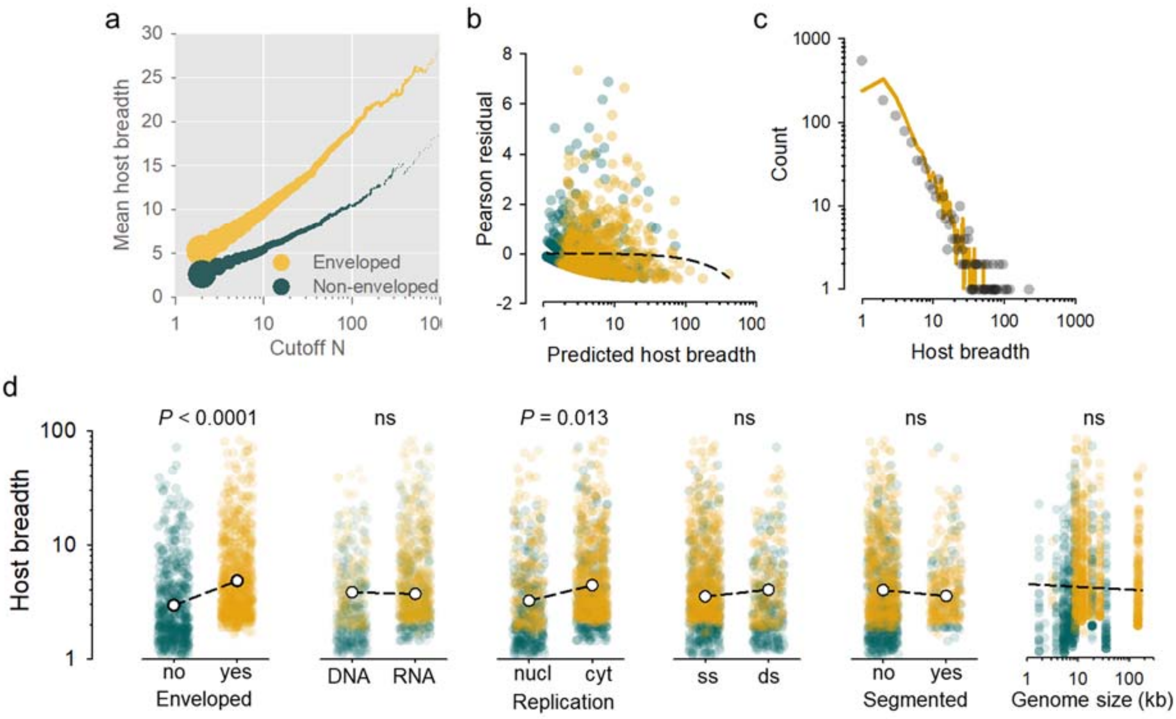


**Figure S4. Analysis of viral host breadth. a.** Viral host breadth increases more rapidly with the cutoff N for enveloped viruses than for non-enveloped viruses. Dot sizes are proportional to the number of viruses included for each cutoff. The dots are shown for $N \geq 2$, $N \geq 3$, and so on. Dots for $N \geq 1$ were too large for visualization and are omitted. **b.** Pearson residuals from a negative binomial regression using a log link function. Orange dots indicate enveloped viruses and green dots non-enveloped viruses. The dashed line shows a least-squares fit to the data using a quadratic function. The model tends to underestimate large host breadths. **c.** Observed host breadths (grey dots) and predicted (orange line) host breadths using the negative binomial regression model. The distribution of observed host breadths was strongly overdispersed, with a mean of 6.40 species, variance equal to 194.13, and a median of 2 species for data satisfying $N \geq 5$. Overdispersion justified the choice of a negative binomial model over a simpler Poisson regression, although a negative binomial did not fully capture the large variance. **d.** Results of the negative binomial regression. For each viral feature considered, scatter plots show the predicted host breadth for each of the 1305 viruses considered. Orange dots indicate enveloped viruses and green dots non-enveloped viruses. White dots and dashed lines indicate the marginal mean predicted by the model. P-values for each predictor variable are shown.
